# Comparative Network Reconstruction using Mixed Integer Programming

**DOI:** 10.1101/243709

**Authors:** Evert Bosdriesz, Anirudh Prahallad, Bertram Klinger, Anja Sieber, Astrid Bosma, René Bernards, Nils Blüthgen, Lodewyk FA Wessels

## Abstract

New anti-cancer drugs that specifically target oncogenes involved in signalling show great clinical promise. However, the effectiveness of such targeted treatments is often hampered by innate or acquired resistance due to feedbacks, crosstalks or network adaptations in response to drug treatment. Addressing this problem requires an understanding of these networks and how they differ between cells with different oncogenic mutations or between sensitive and resistant cells. Here, we present Comparative Network Reconstruction (CNR), a computational method to reconstruct signaling networks based on incomplete perturbation data, and to identify which edges differ quantitatively between two or more signalling networks. Prior knowledge about network topology is not required but can straightforwardly be incorporated. We extensively tested our approach using simulated data and applied it to perturbation data from a BRAF mutant cell line that developed resistance to BRAF inhibition. Comparing the reconstructed networks of sensitive and resistant cells suggests that the resistance mechanism involves re-establishing wildtype MAPK signaling, possibly through an alternative RAF-isoform.

## Introduction

Aberrations in cellular signal-transducing networks are one of the hallmarks of cancer [1], and many new anti-cancer drugs specifically target genes involved in signaling [2]. Unfortunately, the effectiveness of targeted therapies is often limited by unexpected feedbacks or crosstalks [3–5], and resistance typically emerges due to network adaptations that reactivate inhibited pathways [6, 7]. The design of effective treatments can benefit from an accurate and quantitative understanding of the interactions in such networks and how these are remodelled under drug exposure.

Several types of computational approaches have been developed to characterize signaling networks. Ordinary differential equation based models give detailed, dynamic and quantitative descriptions of a system [8]. However, such models typically have many parameters that need to be estimated. Fitting these requires vast amounts of measurements and is computationally very expensive, limiting the scope to small systems of which the topology is already well known. Because of their relative simplicity, logic (or Boolean) models are suitable for larger systems [9], and have been used to predict synergistic drug interactions [10] and to identify differences in signaling network topologies between cell lines [11]. However, these models are not quantitative and require discretization of the data.

Modular response analysis (MRA) is a framework that strikes a balance between the level of detail and scope of a model [12,13]. MRA is a mathematical framework that quantifies the interaction strengths between network nodes based on perturbation data. It only considers interactions between ‘modules’, characterized by a single output affecting other modules. In the context of signaling networks, this means that only the active form of the kinase needs to be considered. MRA is considerably simpler than ODE-based models while retaining the relevant information about how network nodes influence each other. In its original formulation, MRA and similar methodologies [14] require perturbations of all nodes, which is not always feasible, for instance, because not all proteins are druggable. This problem can be solved by a maximum-likelihood [4,15] or Bayesian [16,17] reformulation of MRA, or similar methods [18], but these methods typically require a defined network topology as input.

While all existing approaches model a specific given signaling network, one is often specifically interested in the quantitative *differences* between signaling networks. For instance, how signaling networks in wild type cells are transformed by oncogenic mutations or revealing resistance mechanisms that appear under drug exposure by comparing the treatment-naive and resistant networks. Here, we present Comparative Network Reconstruction (CNR), a reformulation of MRA as a Mixed Integer Quadratic Programming (MIQP) problem that allows for efficient network reconstruction and quantification of networks in multiple cell lines simultaneously, allowing the identification the most relevant differences between them. As input, CNR uses possibly incomplete perturbation data in combination with annotation of the nodes that are directly affected by the perturbations. CNR does not require a predefined network topology, but prior information can be included straightforwardly.

We first extensively test CNR on simulated perturbation data, exploring how noise and incomplete perturbation data affect the quality of network reconstructions. We then demonstrate our approach using perturbation data from a BRAF mutant, PTPN11 KO colorectal cell line that acquired resistance resistance to BRAF inhibition. The resistant cells are less sensitive to BRAF inhibition, more sensitive to growth-factor stimulation, and have a negative feedback from ERK to MEK, suggesting that the resistance mechanism involves re-establishing wild type MAPK signaling.

## Materials and Methods

### Formulation of the Comparative Network Reconstruction algorithm

Here, we briefly describe the Comparative Network Reconstruction formalism. For more background and detail, please consult the Supporting Information.

CNR is based on Modular Response Analysis (MRA) [12], a mathematical framework to identify and quantify the interactions in a network based on perturbation experiments. MRA links the measured response *R_ij_* (defined as the log2-fold change in the steady-state activity of node *x_i_* after perturbation *j*) to the unobserved physical interaction strengths *r_kl_* (defined as the logarithmic partial derivative of *x_k_* with respect to *x_l_*) and direct perturbation effects *s_mn_* (the scaled direct effect of perturbation *n* on *x_m_*). The matrices of local and global response coefficients and the direct perturbation effects are related through the equation

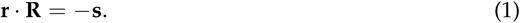

CNR aims to find a network model that solves these equations for multiple cell lines simultaneously while minimizing model complexity and the number of differences between the reconstructed networks of the different cell lines.

Formally, given *N_n_* nodes in the network, *N_p_* perturbations and *N_c_* cell lines, and for *i*, *j* ∈ {1,2,…,*N_n_*}, *n* ∈ {1,2,…,*N_p_*} and *x* ∈ {1,2,…,*N_c_*}, the MIQP problem for reads as follows:

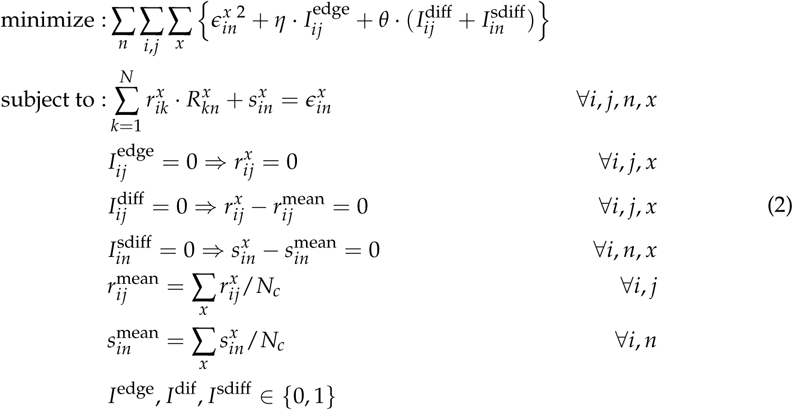

where *η* and *θ* are hyper parameters to tune the degree of penalization on the number of edges and difference between cell lines, respectively. 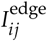, 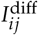 and 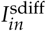 are indicator variables indicating whether an edge is present or not, and whether and edge or pertubation strength is allowed to differ between cell lines, respectively.

Prior information about network topology can be added as constrains of the form 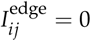 or 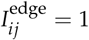 for edges that are known to be absent or present, respectively. Similarly, information about the sign of an edge or perturbation can be added as a constraint of the form 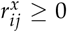 or *s_in_* ≤ 0.

The optimization problem is solved using IBM ILOG CPLEX solver (Version 12.7.1), which is freely available for academic use. Importantly, the MIQP solver guarantees optimal solutions (within small numerical tolerances). In some cases, we also want to obtained close-to-optimal solutions. For this, we use the CPLEX solution-pool functionality. CNR is implemented using a python interface available on https://bitbucket.org/evertbosdriesz/cnr.

### Simulating perturbation data

To generate simulated perturbation data, we used an ODE model of EGFR signaling developed by Orton et al. [19]. We first approximated the steady state by integrating the model over time and subsequently solving the steady state equations (i.e. setting all derivatives equal to 0). Activating mutations in BRAF, RAS and EGFR were modeled by setting all these proteins in the active state in the initial conditions, and setting the conversion rate to the inactive state to zero. We simulated perturbations as knockdowns of a protein by reducing the total amount of that protein by 50%, and recalculating the steady state. We calculated the global response coefficient for protein *i* in response to perturbation p as *R_ip_* = log_2_(*X_ip_*/*X_i_*_,0_), where *X_i,p_* and *X_i_*_,0_ are the perturbed and unperturbed steady state concentrations of the active form of protein *i*, respectively. We added noise to the global response coefficients by multiplying them with a random number drawn from a normal distribution with mean (*μ*) set to 1 and standard deviation (*σ*) set to the noise level.

### Cells and cell culture

All cell lines were derived from the the BRAF^V600E^ mutant VACO432 colorectal cancer cell line. Generation of the VACO432 PTPN11 KO clones is described by Prahallad et al. [20]. Persister cells were generated by prolonged culturing of VACO432 PTPN11 KO cells in the presence of 2 μM Vemurafenib and selecting surviving colonies. Cell lines were were cultured in RPMI supplemented with 10% fetal calf serum (FCS) 1% Glutamine and 1% Penicillin/Streptomycin (Gibco).

### Perturbation experiments

Prior to the perturbation experiments, cells were synchronized by serum starvation during 24 hours. During this period, the persister cells were kept exposed to Vemurafenib. At t = -60 minutes, cells were treated with an inhibitor (BRAF: Vemurafenib at 2 μM, MEK: Selumetinib at 1 μM, ERK: SCH772984 at 1 μM, AKT: MK2206 at 1 μM, or PI3Ki: GDC0941 at 1 μM; All SelleckChem). At t=0, cells were stimulated with growth factor (EGF at 20ng ml^-1^, HGF at 25 ng ml^−1^ or NRG1 at 25 ng ml^−1^; all R&D systems) and at t=30 min were harvested by washing with ice-cold PBS and lysing with with Bio-Plex Pro Cell Signaling Reagent Kit (Bio-Rad) lysis buffer. Cell lysate was analyzed with the Bio-Plex Protein Array system (Bio-Rad, Hercules, CA) according to the suppliers protocol as described previously [4]. The following phospho-sites were measured: EGFR^Y1068^, MEK^S217,S221^, ERK1/2^T202,Y204/T185,Y187^, p90RSK^S380^, RPS6^S235/S236^, PI3K p85^S15^, AKT^S473^, mTOR^S2448^, and GSK3A/B^S21/S9^.

## Results

### An efficient method for network reconstruction from perturbation data

We have developed Comparative Network Reconstruction (CNR), a computational framework to 1) reconstruct and quantify interactions between signaling proteins based on potentially incomplete perturbation experiments; 2) doing so in multiple cell lines simultaneously, and 3) with the possibility to use prior data on the network topology and edge signs (Fig 1). CNR is a reformulation and extension of Modular Response Analysis (MRA) [12] as a mixed integer quadratic programming (MIQP) problem.

**Figure 1:**
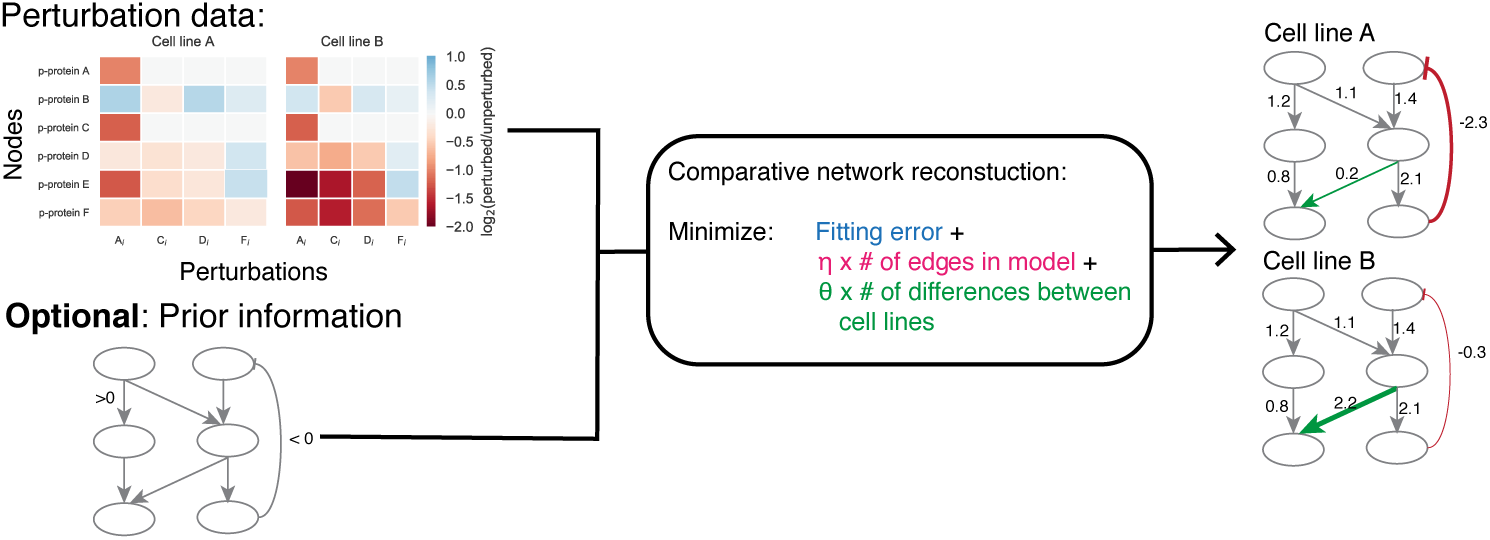
Schematic illustration of Comparative Network Reconstruction. Comparative network reconstruction requires perturbation data (log2-fold changes of node activity compared to a reference state) of multiple cell lines as input. Optionally, information on the network topology or edge signs can be included. Based on this data, an Mixed Integer Quadratic Programming problem is formulated that aims to find a network model that fits the while penalizing the number of edges in the network and the number of quantitative differences between the cell lines. The output is a network quantification and identification of which edges differ between the cell lines. Edge-weights indicate how strongly a target node responds to a change in the activity in the source node.

Reconstruction of a network using MRA requires each node of the network to be perturbed, because otherwise the problem is under-determined. This is typically not feasible, for instance due to cost constraints or simply because not all nodes can be perturbed by e.g. applying a drug. Furthermore, in the classical formulation, it is not possible to simultaneously reconstruct the networks of several cell lines, which complicates their comparison. CNR solves these problems.

CNR takes the log2-fold changes of signaling proteins in response to perturbations as input, together with annotations of which nodes are perturbed (Fig 1). Optionally, information about the presence, absence, or signs of interactions can also be supplied (Fig 1). CNR then tries to find a network model that 1) fits the perturbations data; 2) maintains the same topology for all cell lines; 3) penalizes model complexity (i.e. the number of edges in the network) to prevent over-fitting, and 4) penalizes the number of edges that differ between cell lines to facilitate identification of the most relevant differences. The latter has the additional advantage of reducing the number of parameters, since most edges are described by the same parameter in all cell lines.

The output of CNR is a network quantification for each cell line, in combination with a quantification of the strengths of the effects of perturbations on their direct targets (Fig 1). The interaction strength between a pair of nodes consisting of a source and target node is referred to as the local response coefficient. This coefficient is interpreted as the percentage change in the target node in response to a 1% change in the source node.

### *In-silico* evaluation

To test how well our method works, we simulated perturbation experiments using an ODE-based dynamic model of the EGFR signalling developed by Orton et al. [19] (Fig 2A). Following Orton et al., we also generated BRAF, KRAS or EGFR mutant versions of the model by forcing these proteins to all be in their active state. We simulated network perturbations by ‘knocking down’ all the relevant nodes in the network, one by one, and using the resulting measurements as input to CNR.

**Figure 2:**
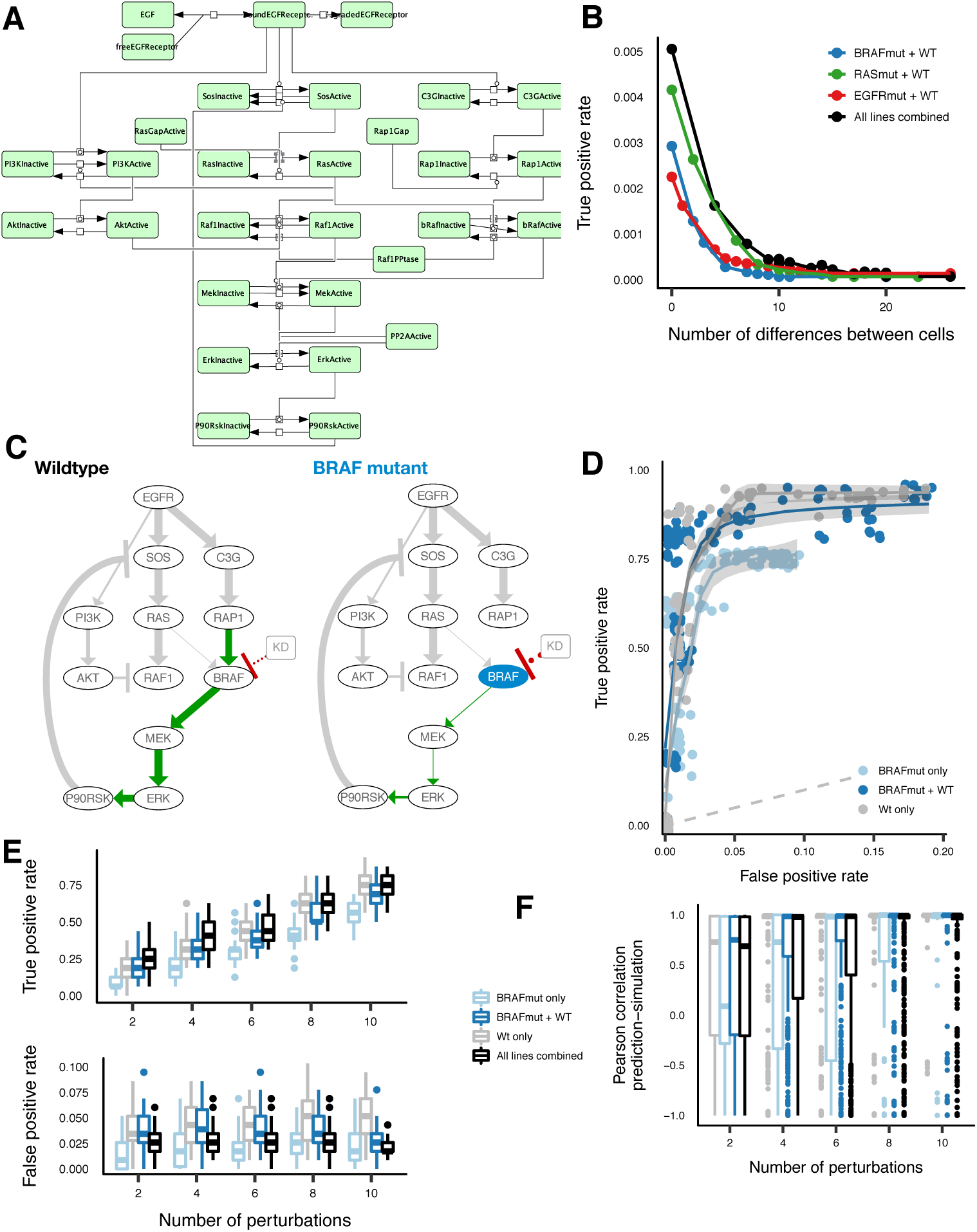
Orton model simulation and CNR results. **A** Schematic of the ODE model used to simulate the perturbation. **B** Fitting error vs number of differing edges between cell lines indicates that most edges can be set to the same value without affecting model fit. **C** Network reconstructions of the wildtype and BRAF mutant model. Edges that differ between the model-reconstructions are highlighted. Green edges are positive, red ones negative. **D** Receiver-operator-curves of network reconstructions using noisy data of the wildtype model, the BRAF model, and both combined. 10% noise was added to the simulated data. **E** True (top) and False (bottom) positive rates of network reconstruction as function of the number of perturbations in the input data. **F** Correlation between predicted and simulated response for perturbations that were not used in reconstructing the network.

We tested the performance of CNR by comparing network reconstructions to the known model from which the data were simulated. Note that since CNR aims to quantify the effect of a perturbation on the steady state concentration of the active form of a protein only, the CNR network reconstruction will be much simpler than the original model. Nevertheless, the true interactions between active proteins are unambiguously defined. Specifically, we wanted to test how the penalty on differences between cell line, noise in the data, or incomplete data affect the ability to reconstruct or quantify the network interactions. In the following sections, unless stated otherwise, we did not employ any information about the model topology for the reconstructions.

#### Effect of the penalty on cell line differences

We first tested how the penalty on the differences between cell lines affects the model fit to the data by performing reconstructions using different values for the *θ*-hyperparameter (using a fixed *η* = 0.005). We first performed this analysis by pairing the wild type with the BRAF, RAS and EGFR mutants, respectively. Then we performed the analysis for all cell lines combined (Fig 2B). For each of these analyses, a model with only a small number of differences (< 10) between the cell lines fits the data nearly as well as a model where all edges differ between the cell lines. The number of edges that differ between cell lines can thus be strongly decreased without affecting the fitting error, which is what one would expect when comparing isogenic cell lines that only differ by a single mutation.

#### Example reconstructions

Figure 2C shows a reconstruction of the wildtype and BRAF mutant models that were reconstructed together (using *η* = 0.005 and *θ* = 0.01). The thickness of the arrows represents the magnitude of the reconstructed local response coefficients. The edges that differ between the cell lines are highlighted in color.As expected, in the BRAF-mutant, BRAF activation is unresponsive to RAP1 and more responsive to knockdown of BRAF itself. Furthermore, all other differences between the cell lines are downstream of BRAF. (In the mutant, the nodes downstream of BRAF are less responsive to changes in their direct upstream kinases due to saturation effects.) Similar results are obtained for the other mutants (Fig S1). This illustrates that a comparative reconstruction can identify where in the network two cell lines differ.

#### Effect of measurement noise

Next, we tested the ability of our method to reconstruct the network topology when the data is noisy. To this end, we performed reconstructions with noise added to the input data, for different values of the *η*-hyperparameter. By starting with a large *η* and gradually lowering it, we get solutions with an increasing number of edges. From these solutions we calculated the true positive rate (number of true positive edges/total number of real edges) and the false positive rate (number of false positive edges/total number of real negative edges). Figure 2D shows the receiver-operating characteristic (ROC) curve for reconstructions of the BRAF mutant (light-blue) and the wild type (gray) cell lines, and simultaneous reconstruction of both together (dark blue, using *θ* = 0.01), when 10% noise is added to the data. A true positive rate of >0.8 is attained while keeping the false positive rate well below 0.05, so despite noisy data, a large majority of the edges can be correctly identified. The performance of the wild type + BRAF-mutant and wild type alone reconstructions are comparable, and better than that of the BRAF-mutant alone. This is presumably because due to constitutive BRAF-activation nodes downstream of BRAF are nearly unresponsive to perturbations upstream of BRAF, making these less informative for identifying downstream edges. Similar results were obtained for the the other mutants (Fig S2). As expected, increasing the noise levels reduces the performance, but even at 100% noise, the reconstruction performs much better than randomly reconstructing the network, represented by the dotted gray line where the true positive rate equals the false positive rate (Fig S2). Interestingly, the performance of the BRAF and the RAS mutant cell line reconstructions suffer less from increasing noise than those of the wild type.

#### Effect of incomplete perturbation data

It is not always feasible to perturb each node in the network. To test how incomplete perturbation data affects the accuracy of network reconstruction, we performed network reconstructions based on a reduced number of input perturbations. To this end, we randomly selected between 2 to 10 nodes to perturb, added 10% noise to the data, and performed the network reconstruction. (Because we expect the total fitting error to scale with the number of perturbations *N_p_*, we scaled *η* and *θ* using *η* = 0.01 * *N_p_/N_n_* and *θ* = 0.01 * *N_p_*/*N_n_*, where *N_n_* = 12.) We repeated this 50 times for each number of perturbations. Figures 2E and S3 show the true and false positive rates of these reconstructions. As expected, the performance decreases with decreasing numbers of perturbations. However, with as few as two or four perturbations, we still attain performance levels well above random guessing. With low numbers of perturbations used as input, there is a clear benefit of combining multiple cell lines, as the panel combining all cell lines consistently has among the highest true-positive and lowest false-positive rates.

#### Predicting the effect of new perturbation

Sometimes one is more interested in predicting the effect of new perturbations from a systems were the topology is known, rather than identifying which edges are present. We therefor next tested how well CNR is capable of predicting the effect of perturbations that were not used in the reconstruction. We again randomly selected between 2 to 10 perturbations, added 10% noise to the data, and performed reconstructions this time with the correct network topology supplied. We used to obtained network-quantifications to predict the response to perturbations that were not used in quantifying the network, and calculated the Pearson correlation between the predicted and true response. In calculating the correlation, we excluded the perturbed node itself since the direct-perturbation effect is not part of the prediction. Perturbations were none of the nodes respond are also excluded. Figures 2F and S4 show that for as little of four or six perturbations, the median pearson correlation of the predictions is very close to 1. Interestingly, while the performance in edge reconstruction of the BRAF and RAS mutants were slightly better than those of the wildtype, their quantitative predictions are slightly worse.

### Network quantification of PTPN11 KO cells with and without acquired resistance to BRAF inhibition

To test the ability of CNR in a real-world setting, we set out to identify the cause of resistance to targeted anti-cancer drugs by elucidating how the signaling network changed in resistant cells. Specifically, we wanted to understand how a PTPN11 KO colorectal cancer (CRC) cells harboring an activating BRAF^V600E^ mutation become resistant to BRAF inhibition with Vemurafenib.

#### PTPN11 KO and sensitivity to BRAF inhibition

Normally, BRAF^V600E^ mutated CRC cells are unresponsive to BRAF inhibition due to a feedback loop that activates the receptor tyrosine kinase (RTK) EGFR upon BRAF inhibition [3, 4]. Recently, Prahallad et al. showed that knockout of PTPN11 – which is required to transduce signals from RTKs to the MAPK pathway – prevents this feedback loop from reactivating the MAPK signaling pathway [20] (Fig 3A). However, when PTPN11 KO cells are cultured in the presence of a BRAF inhibitor, they eventually become resistant to this inhibitor.

**Figure 3:**
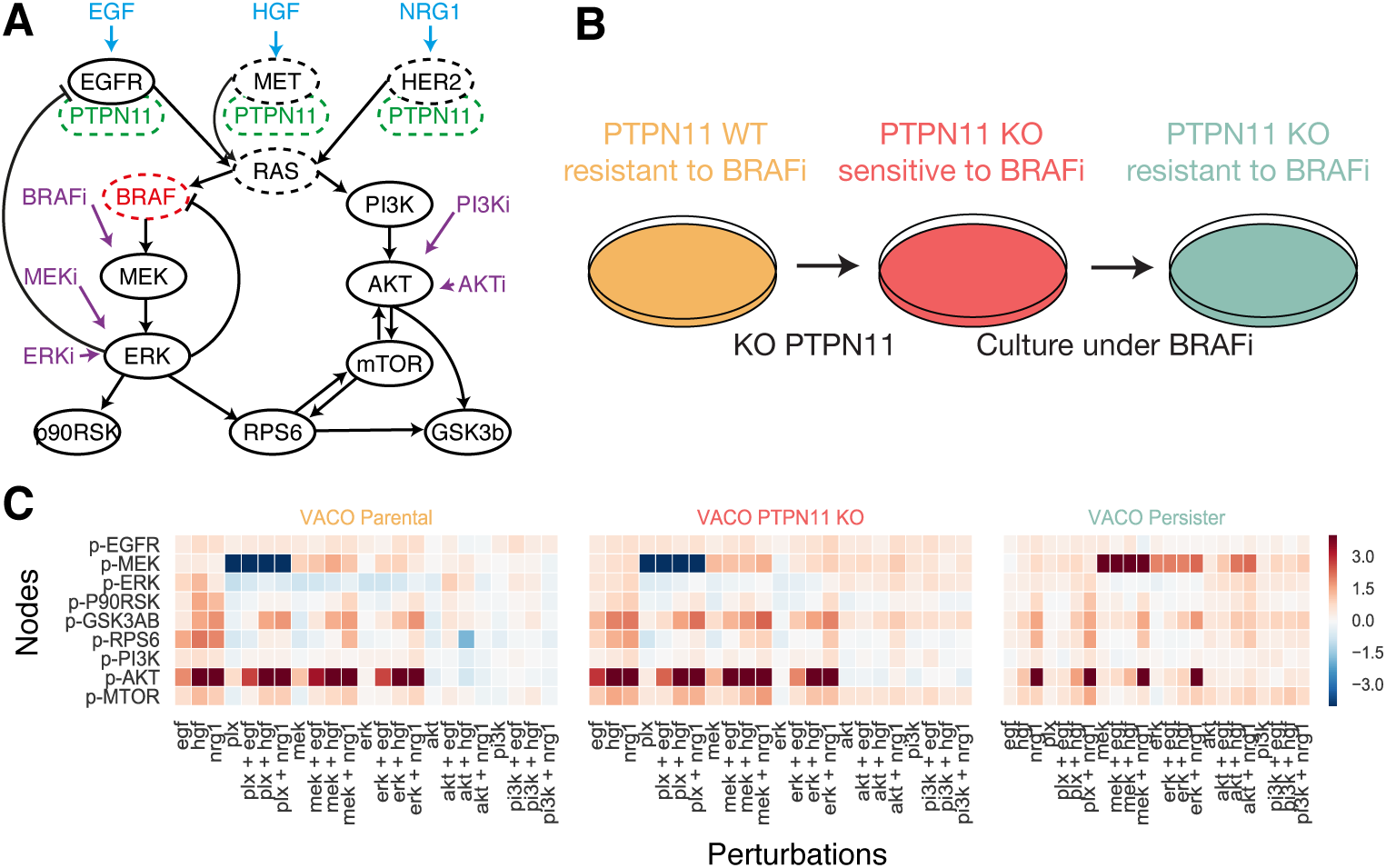
Persister experiments. **A.** Schematic of role of PTPN11 in signal transduction. **B.** Overview of the cell-lines used. VACO432 cells are insensitive to BRAF inhibition, but can be sensitized by knocking out PTPN11. Prolonged culturing VACO432 PTPN11 KO cells in the presence of BRAF inhibitor gives rise to resistant VACO432 PTPN11 KO cells. **C.** Results of the perturbation experiments. The color scale indicates log2-fold change relative to unstimulated, uninhibited controls.

#### Generation of persisters

In order to test how resistance to the BRAF inhibition in PTPN11 KO CRC cells emerges, we generated ‘persister’ cells by culturing a PTPN11 KO clone of the BRAF^V600E^ mutated CRC cell line VACO432 in the presence of the BRAF inhibitor Vemurafenib. After roughly 2 weeks, persister cells emerged that were able to grow in the presence of Vemurafenib at a growth rate comparable to that of the parental line in the absence of drug (Fig 3B).

#### Perturbation experiments

We systematically perturbed VACO432 parental, PTPN11 KO, and the derived persister cells with combinations of growth factors and inhibitors targeting proteins in the PI3K-AKT and MAPK pathway, and measured the steady-state responses of the phosphorylation status of main proteins in the pathways (Fig 3A and C). The log2-fold changes, relative to their unperturbed (i.e. uninhibited an unstimulated) state, were used as input to the CNR optimization problem. As the MAPK and AKT pathways are relatively well characterized, we did not focus on the reconstruction of the pathway topology. Instead, we restrict the model to consist of known interactions and feedbacks obtained from the literature (Fig 3A).

#### Modeling of perturbations

Some kinase inhibitors prevent activation of their target by blocking the phosphorylation of the kinase itself, whereas other work by preventing the kinase activity of the phosphorylated protein. We model the latter type of inhibitor as perturbing the substrates of its direct target. Specifically, the MEK and PI3K inhibitors are preventing the kinase activity of their targets and are thus modeled to perturb ERK and AKT, respectively. Since we did not measure BRAF, we cannot model the direct effect of BRAF inhibition on BRAF, but instead model it as a perturbation of MEK. Similarly, we didn’t measure activity of HER2 and MET, so we model HGF and NRG1 stimulation as affecting MEK, PI3K and AKT directly.

#### Network reconstruction

Figure 4A shows the network reconstruction of the 3 cell lines, and how they differ. We allowed for 6 differences between networks as it gives a good balance between model-fit and model-complexity (Fig S5). As expected, the response of MEK activation to BRAF inhibition is much smaller in the persisters compared to the parental and the PTPN11 KO cells. Furthermore, in the persisters the feedback from ERK to MEK is stronger and MEK is more responsive to receptor stimulation. Finally, the effect of EGFR on AKT is weaker in both the PTPN11 KO and the persister cells, indicating that in PTPN11 might be involved in signaling to the AKT pathway as well as the MAPK pathway.

**Figure 4:**
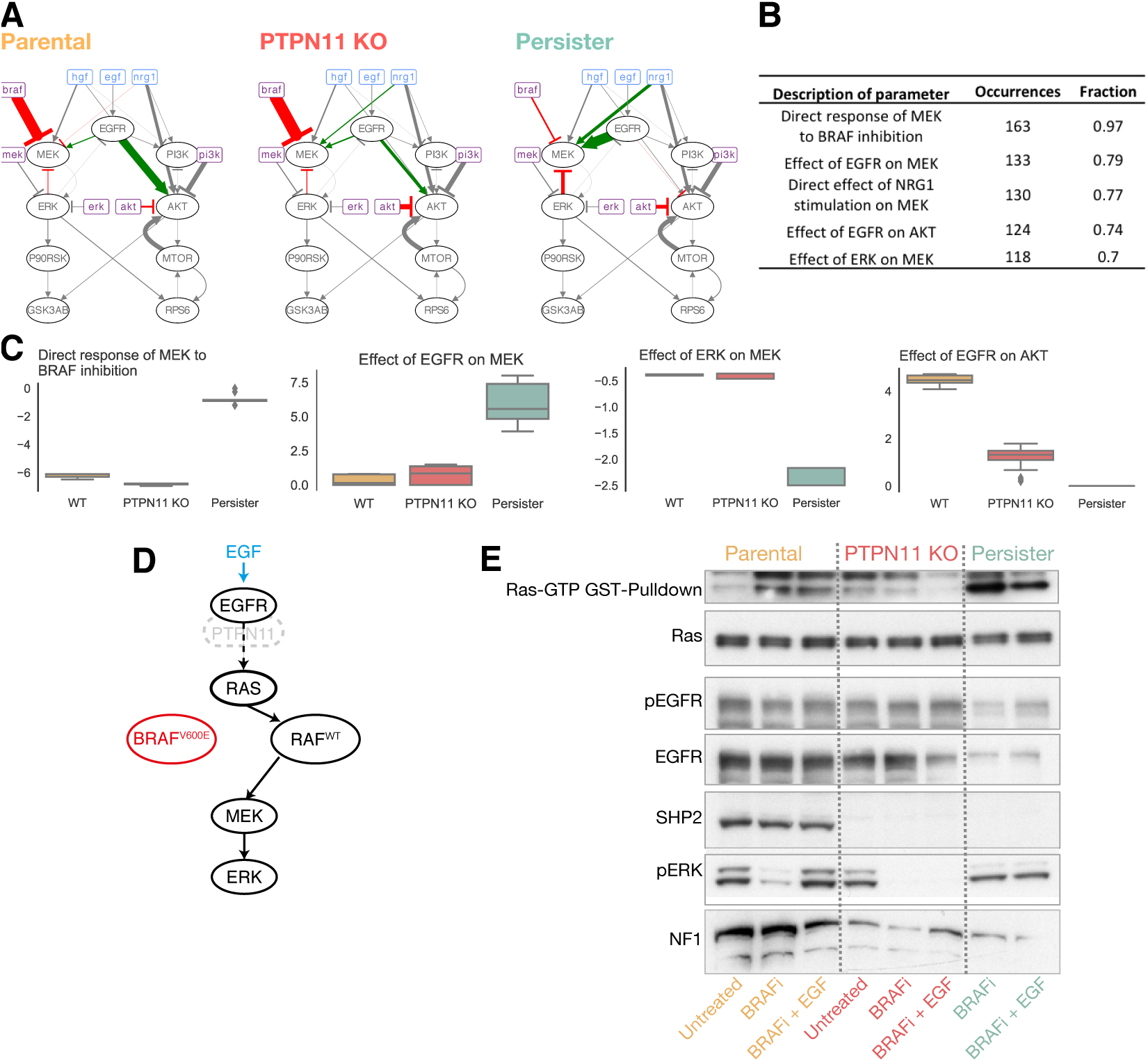
Model reconstruction of sensitive and persister VACO432 cells. **A.** Network reconstruction from perturbation data. **B.** Overview of parameters that differ in many of the solutions. **C.** Boxplots of selected parameters indicating that the differences are quantitatively similar in different solutions. **D.** Hypothesis: Resistance is due to re-establishment of normal MAPK signaling. **E.** Consistently, pERK is activated and GTP constitutively loaded with GTP.

#### Robustness of network reconstruction

To test how robust these results are, we generated a large pool of solutions by varying the allowed number of differences between 0 and 16, and also considering close-to-optimal solution (with an objective value at most 1.5 times the best possible given the number of differences between cell lines). We counted how often each edge differed between cell lines in the 168 solutions in the solution-pool. There is a limited number of edges that occur in the majority of these solutions (Fig 4B). These are: 1) the response of MEK to BRAF inhibition; 2) the effects of EGFR on MEK and AKT; 3) the feedback from ERK to MEK; 4) the responsiveness of MEK to growth-factor stimulation and 5) the responsiveness of AKT to AKT inhibition. Furthermore, the often-occurring differences were also quantitatively consistent in most solutions (Fig 4C), and in agreement with the observations we made earlier: 1) MEK activity in persisters is less responsive to BRAF inhibition; 2) more responsiveness to growth factor stimulation, 3) persisters have a negative feedback from ERK to MEK and, 4) the connection between EGFR and AKT is stronger in the (PTPN11 WT) parental cells. This indicates that these results are robust and do not depend on the detailed settings in the network reconstruction.

#### Interpretation

The CNR results suggest a model in which the persister cells became resistant by re-establishing normal (RAF-wildtype) MAPK signalling, by, for instance, using an alternative RAF-isoform (Fig 4D). Such a model explains the reduced response of MEK to Vemurafenib, which specifically targets BRAF^V600E^ and not wildtype RAF. It explains the stronger negative feedback from ERK to MEK, which is mediated through RAF and thus weak if BRAF is constitutively active. It also explains the increased responsiveness to EGFR and other growth factors, which are also mediated through RAF and thus weak if BRAF is constitutively active. Consistent with this hypothesis, pERK is activated in the persisters and RAS is constitutively loaded with GTP (Fig 4E). This hypothesis suggests that the persister cells might be sensitive to pan-RAF inhibition.

### Discussion and conclusion

In this study we developed Comparative Network Reconstruction (CNR), a computational method to reconstruct signaling networks based on perturbation data. CNR focusses on identifying quantitative differences between a set of cell lines (or other experimental models), as this is often the relevant biological question. One of the main advantages of CNR is it’s flexibility regarding prior knowledge of the network topology. Such knowledge it is not required, but can straightforwardly be incorporated. Using simulated data we showed that CNR works well with noisy and incomplete data.

We applied CNR to perturbation data we generated from BRAF^V600E^ mutated, PTPN11 KO colorectal cancer cells that were made resistant to the BRAF inhibitor Vemurafenib through continuous exposure to sub-lethal doses of the drug. The network reconstructions suggested a reversion to RAF wildtype-like MAPK signaling in the resistant cells that would likely make them sensitive to pan-RAF inhibition. This hypothesis raises a number of other interesting questions, such as how RAF can be activated and responsive to growth factor stimulation in the absence of PTPN11.

CNR is formulated as an Mixed Integer Quadratic Programming (MIQP) problem. Theoretically, such problems are NP-hard, and the search space increases exponentially with the number of nodes in the network. In practice the MIQP problem associated with CNR can typically be efficiently solved. For example, a typical reconstruction for the 12-node Orton model is optimized in under a second on a standard laptop computer. Parameter-setting that lead to highly connected networks are somewhat slower (Fig S6), but since most proteins interact with a limited set of other proteins, biological networks are sparse and the highly connected solutions are not relevant. Additionally, the run time can be strongly reduced by incorporating prior information; for instance by providing a starting network with well known interactions and allowing for a limited number of additional edges. This approach is thus scalable to much larger networks.

In this work we focused on signaling networks, but the method can be applied to any perturbation data set were the targets of the perturbations are known. We used a L0-penalty for the differences between networks (an edge is either different or not) because such results are convenient to interpret as only a subset of edges need to be inspected. However, the penalty can be replaced with an L1 or L2-penalty (The absolute or square of the difference between two edges, respectively), which in some cases better resembles the biological problem, and is expected be be faster to solve.

To conclude, CNR provides a flexible and efficient way to reconstruct networks from perturbation data. Using simulations, we demonstrated its ability to accurately reconstruct networks when data is noisy and incomplete. Application to a real-world problem showed its robustness and potential in formulating hypotheses on biological problem.

## Supporting Information

### Basics of Modular Response Analysis

Consider a network consisting of *N* nodes, the state of which is specified by **x** = {*x*_1_,…, *x_N_*}. These nodes can be the proteins involved in a signaling network, in which case their state is the concentration of phosphorylated protein. More generally, these nodes are modules of arbitrary complexity, which may involve many cellular components connected by intra-modular reaction. However, by assumption only a single element of the module serves as output. The steady state of the system is defined by:

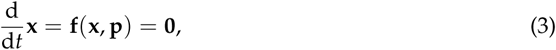

where **p** is vector of parameters. Modular Response Analysis (MRA) aims to reconstruct an interaction network based on measuring the steady state response to (many) perturbations.

A perturbation in *p_j_* can affect *x_i_* directly, but also indirectly trough network effects. Upon a infinitesimal perturbation in *p_j_*, the following relation holds for each node:

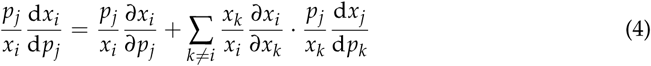

Hence, an experiment of *M* perturbations yield *M* × *N* relations.

In MRA, all terms in equation 4 have a name and an interpretation. The global response coefficient

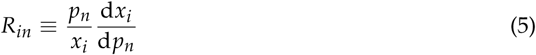

is the response of the node *i* to a perturbation in a parameter *n*. This is typically the quantity that can be measured. The local response coefficient

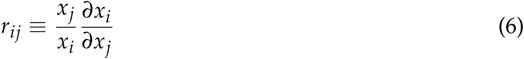

represents the direct effect of a change in *x_j_* on on *x_i_.* By definition, *r_ii_* ≡ −1. The elements of the perturbation matrix

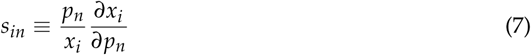

quantify the direct response of node *i* to a perturbation in *p_j_*.

Equation 4 can be reformulated in matrix notation to yield:

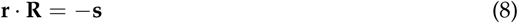

where **r** is an *N* × *N* matrix (with the local response coefficients as its entries) and **R** and **p** are a *N* × *M* matrices. Often, **r** and (and to a lesser extend **s**) are the quantities of interest because they represent the direct interaction. However, these but can not be measured directly because a node cannot be manipulated in isolation. The goal of MRA then, is to find these based on the measured **R**.

The idea behind classical MRA is to perturb each node independently, meaning all nodes are perturbed and that each perturbation affect only one node. In this case, it is possible to select a parameterization such that *s_in_* = *δ_in_* (where *δ* is the Kronecker delta). The solution to the problem than simply is:

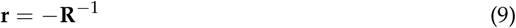

### Formulation of Comparative Network Reconstruction

For the purpose of comparing cell lines (or other experimental models), there are two main problems with MRA. The first is that is typically not feasible to perturb all nodes in a network, which makes the problem under-determined. The second is that it requires fitting a new model for each cell line, which prevents ‘sharing’ data between cell lines and thus might incorrectly identify differences that are the result of e.g. measurement error.

Our strategy to solve this is to reformulate MRA as a Mixed Integer Quadratic Programming (MIQP) problem. MIQP is a form of constrained, non-linear optimization where the objective function is a convex, quadratic function while the constraints are linear, and some variables are integers.

The idea is to look for set of models –one for each cell line-that:

- fits the perturbation data while
- adding a sparsity constraint by penalizing the number of edges in the network, and
- penalizing edges that differ between cell lines.

The fit is found by minimizing **r · R** + **s**. The penalties on the number of, and differences between, edges are implemented using so-called indicator constraints. These are binary variables that can code logical relationship. (Because these indicator constraints are integers, this is a MIQP problem instead of simply a QP one.) For each potential edge *r_ij_* in the network we add to two indicator, 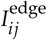 coding for the presence/absence of the edge, and 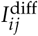, whether it is allowed to differ between cell lines. If 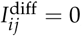, the value of corresponding local response coefficient *r_ij_* is constrained to 0 in all cell lines. Similarly, if l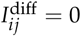, *r_ij_* is constrained to have the same value in all cell lines. The penalties are implemented by adding (weighted) sums over the indicator constraints to the objective functions.

Formally, the MIQP problem reads as follows:

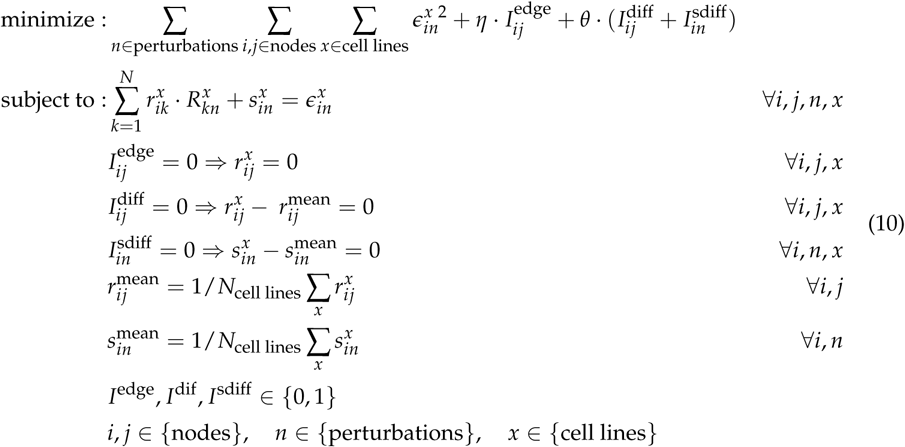

where *η* and *θ* are hyper-parameters to tune the degree of penalization of on the number of edges and difference between edges, respectively.

An important benefit of formulating MRA as an MIQP problem is that prior knowledge can be incorporated straightforwardly by additional constraints.

- Prior information about the network topology can be incorporated by adding constraints on the *I*^edge^. For instance, addition of 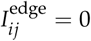 to equation (2) indicates that there is no interaction between node *i* and *j*.
- Prior information on the sign of an interaction can be incorporated by adding constraints on local response coefficients. For instance, if a feedback from node *j* to node *i* is known to be negative, this is encodes by adding 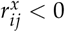 to equation (2).
- Prior information about edges that differ between

Finally, a maximum on total number of edges or differences between edges can be added by simply adding a constraint of the form 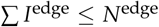 and 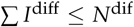.

## Supporting figures

**Figure S1:**
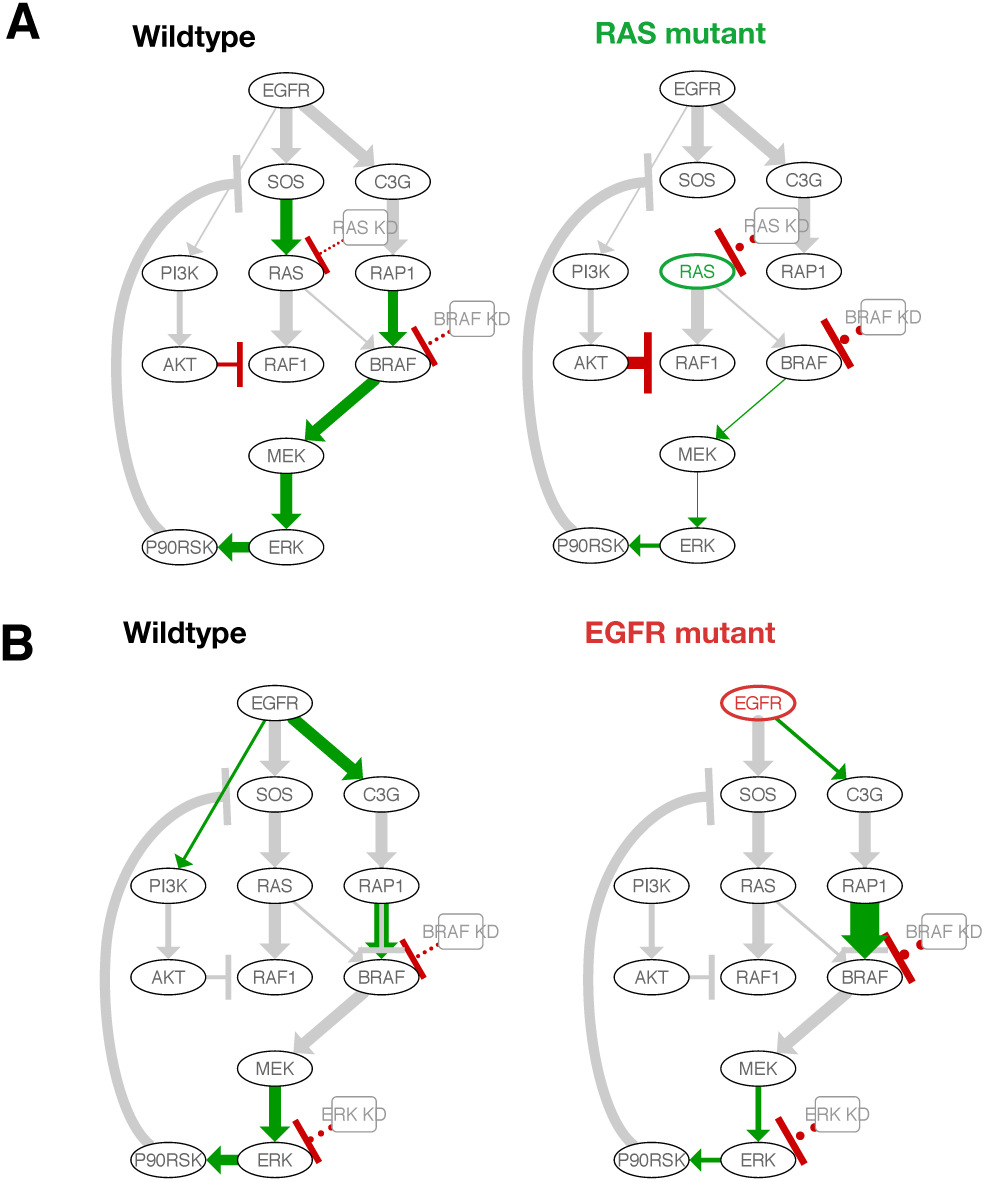
Network reconstruction of RAS-WT and EGFR-WT cell line panels.

**Figure S2:**
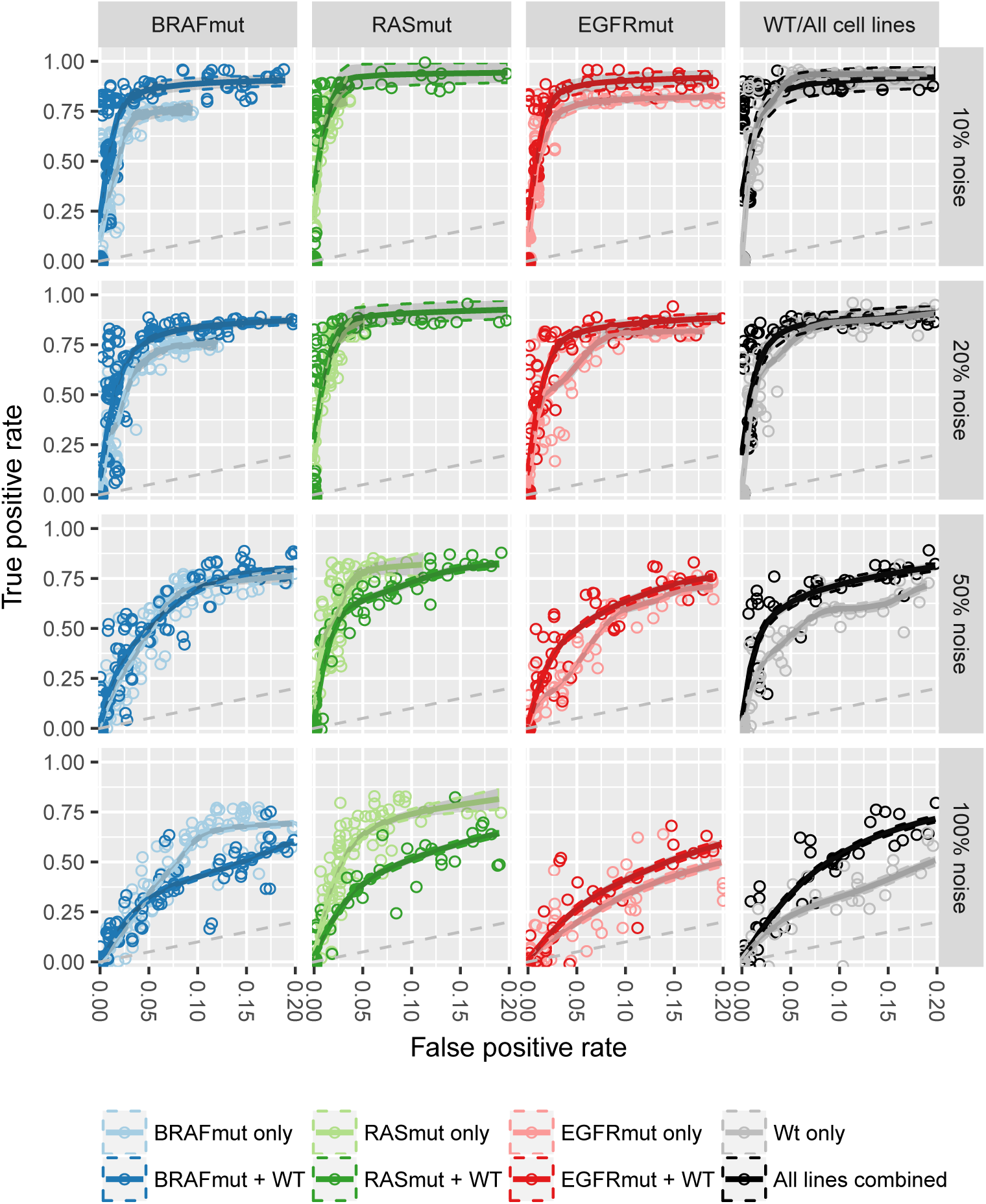
Receiver-operator curves of different cell-line panels with different levels of noise added to the data. By scanning over the *η*-hyperparameter, reconstructions with different numbers of edges were obtained, leading to different true and false positive rates. Reconstructions were performed on either a mutant cell line alone, or together with the wild-type cell line. Additionally, reconstructions on all cell lines simultaneously was performed, and compared to reconstruction on the wild-type cell-line alone. For each noise-level, the scan over the *η*-hyperparameter was repeated 10 times. For each reconstruction, the *θ*-hyperparameter, penalizing the differences between cell lines, was set to 0.001. Each points represents one network reconstruction. Solid lines are estimated by fitting an smooth, monotonically increasing curve through the points using the R scam package [21]. The shaded regions indicate standard error of the means. Note that the false positive rate is only shown until 0.2. For visualization purposes, the points were slightly jittered.

**Figure S3:**
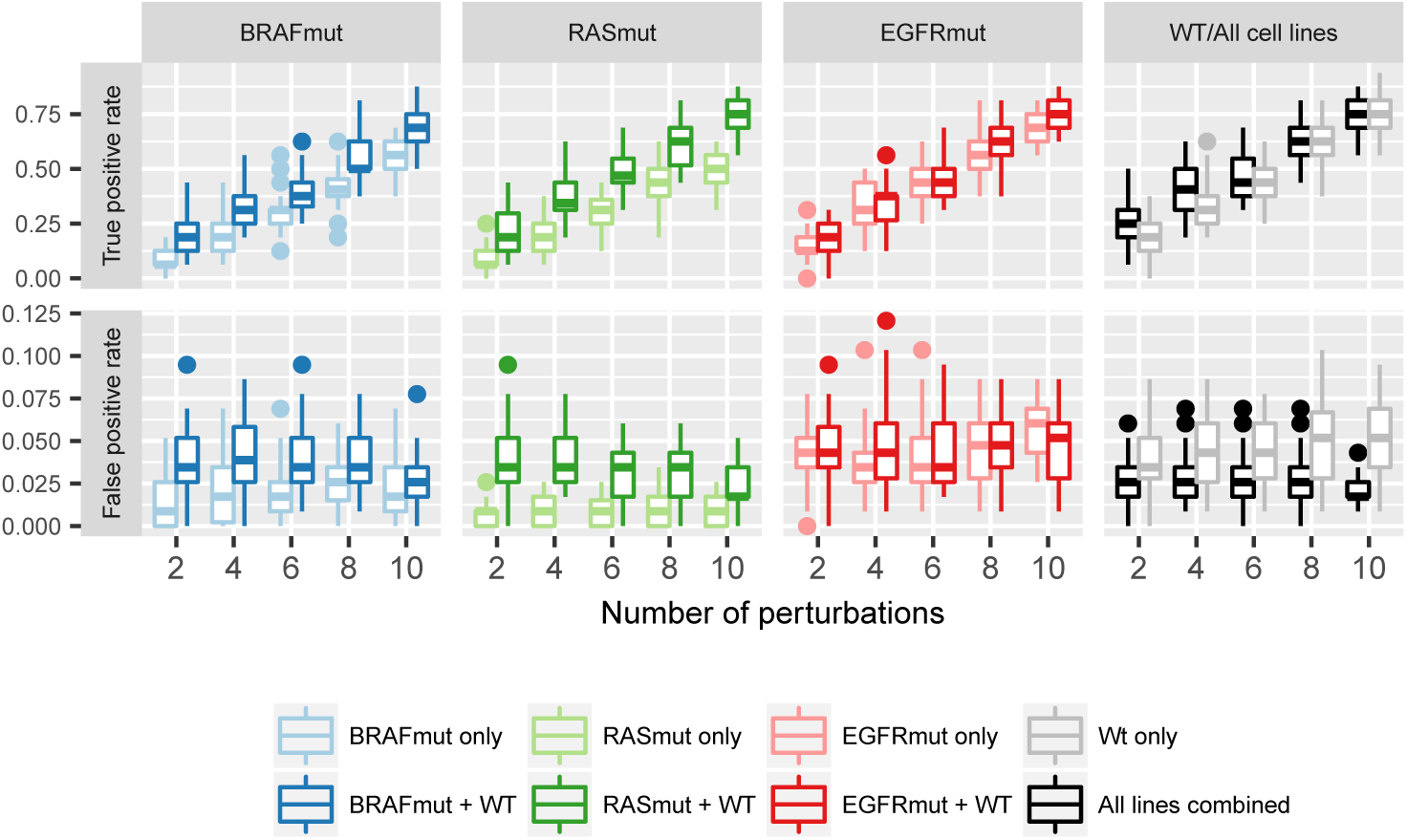
True and false positive rates of edge prediction for different number of perturbations uses for network reconstruction. Reconstructions were performed using different numbers of randomly selected perturbations. For each reconstruction, 10% noise was added to the original global response matrix and a fixed eta and theta were used. To ensure a fair comparison, the same perturbations and noisy global response matrices were used for each panel. The procedure 10 times for each number of different perturbations.

**Figure S4:**
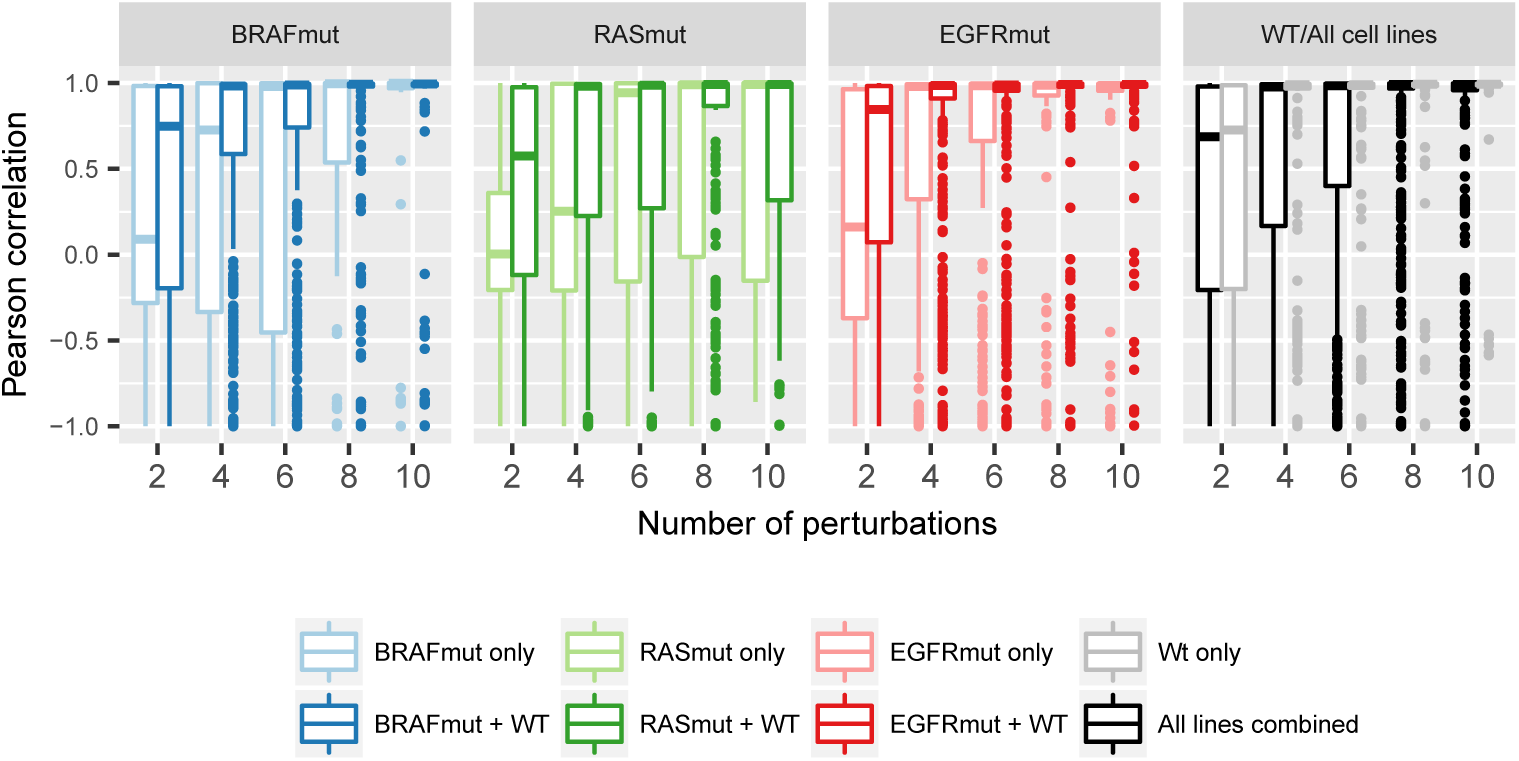
Pearson correlation between predicted and simulated response for different numbers of perturbations used in network reconstruction. Network quantifications were performed using different number of randomly selected perturbations. The correct network topology was provided as input. For each reconstruction, 10% noise was added to the original global response matrix and a fixed eta and theta were used. The obtained network quantifications were used to predict the response to perturbations not used in the reconstruction. The y-axis shows the Pearson-correlation between predicted and true response. For each perturbation, the perturbed node is excluded from the correlations since the direct-perturbation effect is not part of the prediction. Perturbations were none of the nodes respond are also excluded.

**Figure S5:**
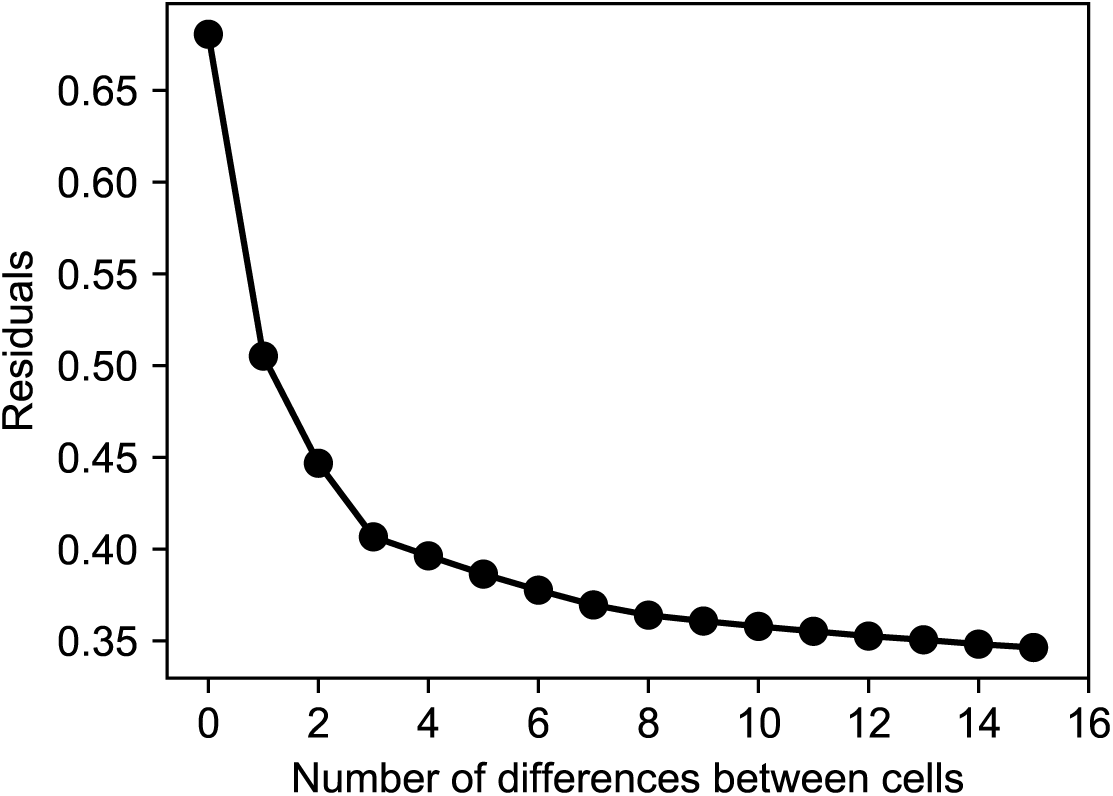
Relation between fitting error and allowed number of differences between cell lines for the VACO432 cell line network reconstructions.

**Figure S6:**
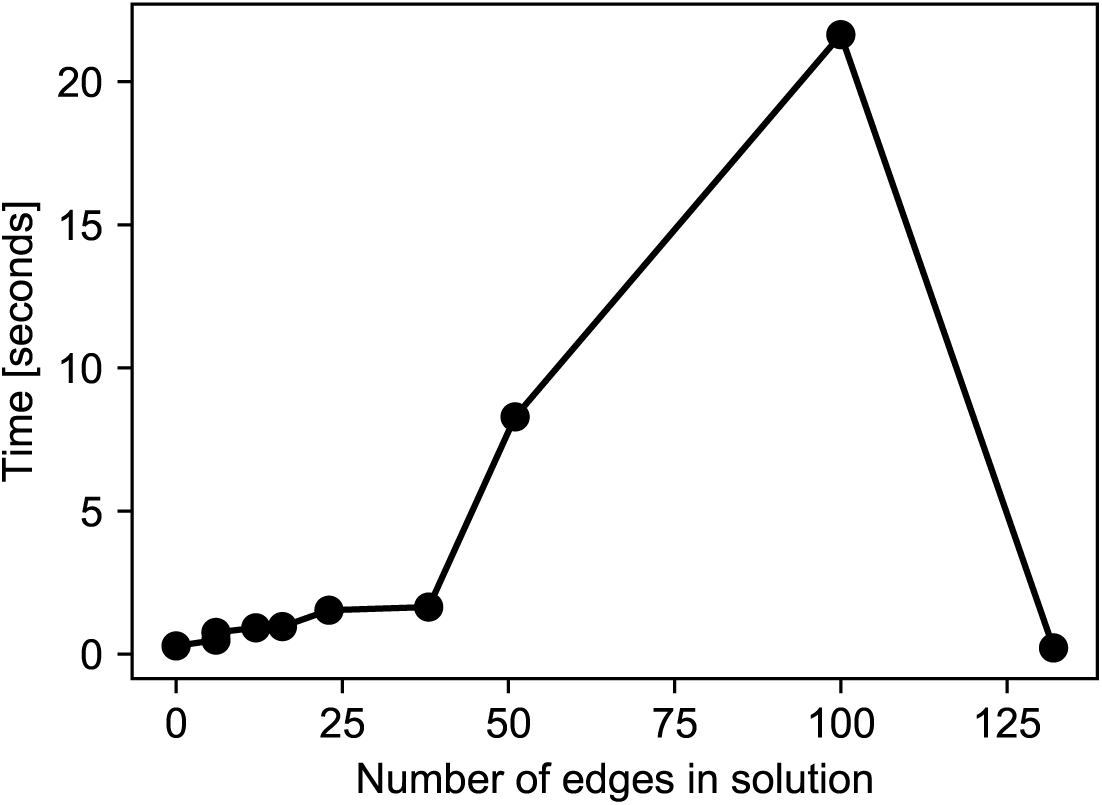
Time to solve the CNR problem as function of the number of edges in the solutions. Denser solutions cost more time. However, as the network becomes almost fully connected, the optimization time starts to decrease because the search-space decreases.

## References

1. Hanahan, D. & Weinberg, R. A. Hallmarks of cancer: the next generation. cell 144, 646–674 (2011).

2. Roberts, P. J. & Der, C. J. Targeting the Raf-MEK-ERK mitogen-activated protein kinase cascade for the treatment of cancer. Oncogene 26, 3291–3310 (May 2007).

3. Prahallad, A. et al. Unresponsiveness of colon cancer to BRAF(V600E) inhibition through feedback activation of EGFR. Nature 483, 100–3 (Mar. 2012).

4. Klinger, B. et al. Network quantification of EGFR signaling unveils potential for targeted combination therapy. Molecular systems biology 9 (Jan. 2013).

5. Sun, C. et al. Intrinsic resistance to MEK inhibition in KRAS mutant lung and colon cancer through transcriptional induction of ERBB3. Cell reports 7, 86–93 (2014).

6. Sun, C. et al. Reversible and adaptive resistance to BRAF(V600E) inhibition in melanoma. Nature (Mar. 2014).

7. Ahronian, L. G. et al. Clinical acquired resistance to RAF inhibitor combinations in BRAF-mutant colorectal cancer through MAPK pathway alterations. Cancer discovery 5, 358–367 (2015).

8. Fey, D. et al. Signaling pathway models as biomarkers: Patient-specific simulations of JNK activity predict the survival of neuroblastoma patients. Science Signaling 8, ra130–ra130 (2015).

9. Saez-Rodriguez, J. et al. Discrete logic modelling as a means to link protein signalling networks with functional analysis of mammalian signal transduction. Mol. Syst. Biol. 5, 331 (Jan. 2009).

10. Flobak, Å. et al. Discovery of drug synergies in gastric cancer cells predicted by logical modeling. PLoS computational biology 11, e1004426 (2015).

11. Saez-Rodriguez, J. et al. Comparing signaling networks between normal and transformed hepatocytes using discrete logical models. Cancer Res. 71, 5400–11 (Aug. 2011).

12. Kholodenko, B. N. et al. Untangling the wires: a strategy to trace functional interactions in signaling and gene networks. Proceedings of the National Academy of Sciences of the United States of America 99, 12841–6 (Oct. 2002).

13. Bruggeman, F. J., Westerhoff, H. V., Hoek, J. B. & Kholodenko, B. N. Modular Response Analysis of Cellular Regulatory Networks. Journal of Theoretical Biology 218, 507–520 (Oct. 2002).

14. Gardner, T. S., di Bernardo, D., Lorenz, D. & Collins, J. J. Inferring genetic networks and identifying compound mode of action via expression profiling. Science 301, 102–5 (July 2003).

15. Stelniec-Klotz, I. et al. Reverse engineering a hierarchical regulatory network downstream of oncogenic KRAS. Molecular Systems Biology 8 (2012).

16. Santra, T., Kolch, W. & Kholodenko, B. N. Integrating Bayesian variable selection with Modular Response Analysis to infer biochemical network topology. BMC Syst. Biol. 7 (2013).

17. Halasz, M., Kholodenko, B. N., Kolch, W. & Santra, T. Integrating network reconstruction with mechanistic modelling to predict cancer therapy. Science signaling 9, ra114 (2016).

18. Korkut, A. et al. Perturbation biology nominates upstream-downstream drug combinations in RAF inhibitor resistant melanoma cells. Elife 4, 1–31 (2015).

19. Orton, R. J. et al. Computational modelling of cancerous mutations in the EGFR/ERK signalling pathway. BMC systems biology 3, 100 (2009).

20. Prahallad, A. et al. PTPN11 Is a Central Node in Intrinsic and Acquired Resistance to Targeted Cancer Drugs. Cell Reports, 1–8 (2015).

21. Pya, N. & Wood, S. N. Shape constrained additive models. Statistics and Computing 25, 543–559 (May 2015).

